# Mucin-induced surface dispersal of *Staphylococcus aureus* and *Staphylococcus epidermidis* via quorum sensing-dependent and independent mechanisms

**DOI:** 10.1101/2024.05.24.595766

**Authors:** Kristin M. Jacob, Santiago Hernandez-Villamizar, Neal D. Hammer, Gemma Reguera

**Affiliations:** Department of Microbiology, Genetics & Immunology, Michigan State University, East Lansing, MI 48824

**Keywords:** staphylococcus, mucin, mucosa, motility, dispersal, dendritic formation, quorum sensing, reservoirs

## Abstract

Nasopharyngeal carriage of staphylococci spreads potentially pathogenic strains into (peri)oral regions and increases the chance of cross-infections. Some laboratory strains can also move rapidly on hydrated agar surfaces, but the biological relevance of these observations is not clear. Using soft-agar (0.3% w/v) plate assays, we demonstrate the rapid surface dispersal of (peri)oral isolates of *Staphylococcus aureus* and *Staphylococcus epidermidis* and closely related laboratory strains in the presence of mucin glycoproteins. Mucin-induced dispersal was a stepwise process initiated by the passive spreading of the growing colonies followed by their rapid branching (dendrites) from the colony edge. Although most spreading strains used mucin as a growth substrate, dispersal was primarily dependent on the lubricating and hydrating properties of the mucins. Using *S. aureus* JE2 as a genetically tractable representative, we demonstrate that mucin-induced dendritic dispersal, but not colony spreading, is facilitated by the secretion of surfactant-active Phenol-Soluble Modulins (PSMs) in a process regulated by the *agr* quorum sensing system. Furthermore, the dendritic dispersal of *S. aureus* JE2 colonies was further stimulated in the presence of surfactant-active supernatants recovered from the most robust (peri)oral spreaders of *S. aureus* and *S. epidermidis*. These findings suggest complementary roles for lubricating mucins and staphylococcal PSMs in the active dispersal of potentially pathogenic strains from perioral to respiratory mucosae, where gel-forming, hydrating mucins abound. They also highlight the impact that interspecies interactions have on the co-dispersal of *S. aureus* with other perioral bacteria, heightening the risk of polymicrobial infections and the severity of the clinical outcomes.

**Importance:** Despite lacking classical motility machinery, nasopharyngeal staphylococci spread rapidly in (peri)oral and respiratory mucosa and cause cross-infections. We describe laboratory conditions for the reproducible study of staphylococcal dispersal on mucosa-like surfaces and the identification of two dispersal stages (colony spreading and dendritic expansion) stimulated by mucin glycoproteins. The mucin type mattered, as dispersal required the surfactant activity and hydration provided by some mucin glycoproteins. While colony spreading was a passive mode of dispersal lubricated by the mucins, the more rapid and invasive form of dendritic expansion of *Staphylococcus aureus* and *Staphylococcus epidermidis* required additional lubrication by surfactant-active peptides (phenol-soluble modulins) secreted at high cell densities through quorum sensing. These results highlight a hitherto unknown role for gel-forming mucins in the dispersal of staphylococcal strains associated with cross-infections and point at perioral regions as overlooked sources of carriage and infection by staphylococci.

## Introduction

Staphylococci are common residents of the nasal and skin microbiota (1) yet readily disperse into the neighboring oral cavity and airways via the pharynx (2). As a result, they are frequently isolated from oral and perioral regions (2–5). Isolates recovered from (peri)oral mucosae include multi-drug resistant strains of *Staphylococcus aureus* and skin commensals such as *Staphylococcus epidermidis* and *Staphylococcus hominis* (4, 5). The presence in these mucosae of *S. aureus* and *S. epidermidis* associated with cross-infections makes the perioral regions overlooked sources of carriage and infection by staphylococci. The lack of machinery (e.g., pili and flagella) for active motility by these bacteria prevents dispersal via swimming as well as surface twitching or swarming. Attachment to and co-dispersal with flagellated motile bacteria such as *Pseudomonas aeruginosa* is however possible (6). Laboratory studies suggest however that some strains of *S. aureus* and *S. epidermidis* may be able disperse on highly hydrated surfaces (7). However, little is known about the biological relevance of these behaviors and whether active cellular processes enable dispersal control.

Some strains of *S. aureus* can passively spread their colonies on soft-agar surfaces provided the medium is sufficiently hydrated (8). Highly hydrated agar media (0.24% w/v agar, 20 min drying time), for example, promoted the passive spreading of many *S. aureus* strains at rates up to 100 µm/min after spot inoculation at high cell densities (8). The spreading phenotype (with or without branched arms or dendrites) and colony expansion rate were strain-specific and dependent on the synthesis and chemical modification of teichoic acids of the staphylococcal cell surface (8). Small reductions in media hydration, such as modest increases in the drying time over 20 min, abolished colony spreading, consistent with a passive mode of dispersal mediated by water carriage (8). Similar colony phenotypes have also been reported in plate assays at higher agar concentrations (0.34% w/v) but shorter (10 min) drying times (9). Under these conditions, some *S. aureus* strains dispersed from the colony edge as slimeless trails (“comets”) that eventually grew into dendrite-like branches (9). These results further supported the notion that water in the medium reduced surface friction and enabled the outward spreading of cells from the growing colonies. Active forms of dispersal have also been proposed, particularly at the high cell densities that trigger the secretion of surfactant-active and lubricating phenol-soluble modulins (PSMs) (8–10). PSM production in *S. aureus* is regulated by the quorum sensing Agr two-component system (AgrC histidine kinase and AgrA response regulator) (11). The phosphorylation of the AgrA response regulator at high cell densities activates the synthesis and secretion of PSMs (7, 8, 11), including two peptides (PSMα3 and PSMγ) with the surfactant activity required for the lubrication of cells moving on semisolid surfaces. A close relative of *S. aureus, S. epidermidis*, has a distinct *agr* system for quorum sensing regulation of PSMs and can also spread on wet surfaces via a mechanism called “darting”, albeit at a much lower speed (6 µm/min) than *S. aureus* (12). While *S. aureus* colony spreading requires lubrication by water in the medium to reduce frictional forces between the cells and the underlying surface, *S. epidermidis* darting results from “the ejection of cells from a capsulated aggregate” (12). *S. aureus* aggregates can also detach and roll away from microcolonies, as if darting, but only under high fluid shear forces (13). In all cases, high cell densities are needed to promote the dispersal of the staphylococcal cells on wet surfaces. The skin commensal *S. hominis* also encodes *agr* quorum sensing systems but lacks most genes for PSM synthesis (14). The laboratory strain AH5009 for example, only produces detectable levels of PSMβ1 and does so under some laboratory conditions (14). To the best of our knowledge, these strains have never been reported to move on surfaces.

The correlation between media hydration, secretion of lubricating PSMs, and staphylococcal movement on semisolid agar surfaces motivated us to study the role that surfactant active, hydrating mucins could have in the mucosal dispersal of staphylococci, particular those inhabiting (peri)oral regions. Human mucosae are typically covered with a mucus layer 10^4^−10^6^ times more viscous than water (15). The controlled secretion of different mucin glycoproteins by the underlying epithelium modulates the hydration, lubrication and viscoelastic properties of the mucus layer across body sites (16). Highly hydrating, gel-forming mucins such as MUC5AC and MUC5B are particularly abundant in the (peri)oral and adjacent respiratory mucus layer (17), making these mucosa a suitable medium for the surface dispersal of potentially pathogenic perioral bacteria into the lower airways. Flagellated bacteria such as *P. aeruginosa* swim within the hydrated media of motility agar (0.3% w/v) plates but cannot move on their surface (swarming) unless the agar concentration is increased to 0.5% (w/v) (18). However, the flagellated cells leverage the presence of as little as 0.05% (w/v) of mucin glycoproteins in the motility agar plates to translocate (“surf”) on the soft-agar surface (18, 19). Surfing motility in *P. aeruginosa* is regulated by quorum sensing, but it is independent of cell density in other surfing species (19). These differences in cell-density control are consistent with a convergent adaptive response for surface dispersal facilitated by mucin lubrication (19). To test for a similar effect of mucins on the dispersal of (peri)oral staphylococci, we investigated the dispersal phenotypes of staphylococci previously isolated by us from the (peri)oral mucosa of healthy young adults (4, 5) and compared the behaviors to those displayed by close relatives among laboratory strains. We describe plate assays for the dissection of passive (outward colony spreading) and active (branching or dendrite formation) dispersal modes induced by surfactant-active, hydrating mucins. Results from this study highlight the stimulatory effect that lubricating mucins have on the spreading of *S. aureus* and *S. epidermidis* colonies, even at low cell densities, and the key role that the *agr*-quorum sensing system plays in modulating PSM secretion to accelerate dendritic expansion at high cell densities. This, and the widespread ability of staphylococcal spreaders to use mucin as a carbon and energy source, confer on these staphylococcal species a competitive advantage for mucosal dispersal in perioral regions and colonization of human airways, where they may cause cross-infections.

## Results

### Mucin induces colony spreading and dendrite formation in (peri)oral isolates of *S. aureus* and *S. epidermidis*

We used soft-agar plate assays typically used to study active motility via swimming (0.3% w/v agar) and swarming (0.5% w/v agar) (20) to investigate the effect of mucin on staphylococcal movement on surfaces (**Fig.1A**). For these experiments, we used highly pure porcine gastric mucin extracted from pig intestines, as described elsewhere (21), and carefully controlled the plate drying time (30 min at room temperature) to ensure reproducibility. Next, we tested nine staphylococcal strains we previously isolated from the oral cavity and perioral regions of four healthy young adults (4, 5). The genomes of these isolates were sequenced with the Illumina platform to retrieve full-length 16S rRNA sequences (Supplemental Table S1) for phylogenetic analysis, which placed the strains as close relatives (bootstrap value >90%) of *S. aureus, S. epidermidis*, or *S. hominis* (**Fig. S1**). None of the isolates spread significantly on plates solidified with 0.5% w/v agar, an assay typically used to study surface motility (swarming) by flagellated species (20) (**Fig. 1A**). Reducing the agar concentration to 0.3% (w/v) to increase the hydration of the medium (swimming motility agar plates) did not stimulate surface spreading of the colonies either (**Fig. 1A**). Furthermore, the colonies had smooth edges and lacked any observable slimeless trails of cells (comets) or branched structures (dendrites) previously reported for some strains of *S. aureus* under similarly hydrated soft-agar conditions (9). However, addition of mucin (0.4% w/v) to these plates stimulated rapid surface spreading of the *S. epidermidis* and *S. aureus* colonies after overnight (18 h) incubation at 37°C (**Fig. 1B**). Additionally, most of the spreading colonies of *S. epidermidis* and *S. aureus* displayed branching structures in their periphery (dendrites) and/or had undulated edges, consistent with emerging and/or growth-consolidated areas of dendritic expansion (**Fig. 1A**). By contrast, the two *S. hominis* strains did not show a mucin-dependent response at 18 h (**Fig. 1**). Longer incubation times (42 h) had only modest effects on the spreading of the *S. hominis* colonies but led to further expansion of the *S. epidermidis* and *S. aureus* colonies (**Fig. 1B**). Hence, mucin-induced dispersal under these laboratory conditions was species-specific, with *S. aureus* and *S. epidermidis* showing the most robust responses.

**Fig 1:**
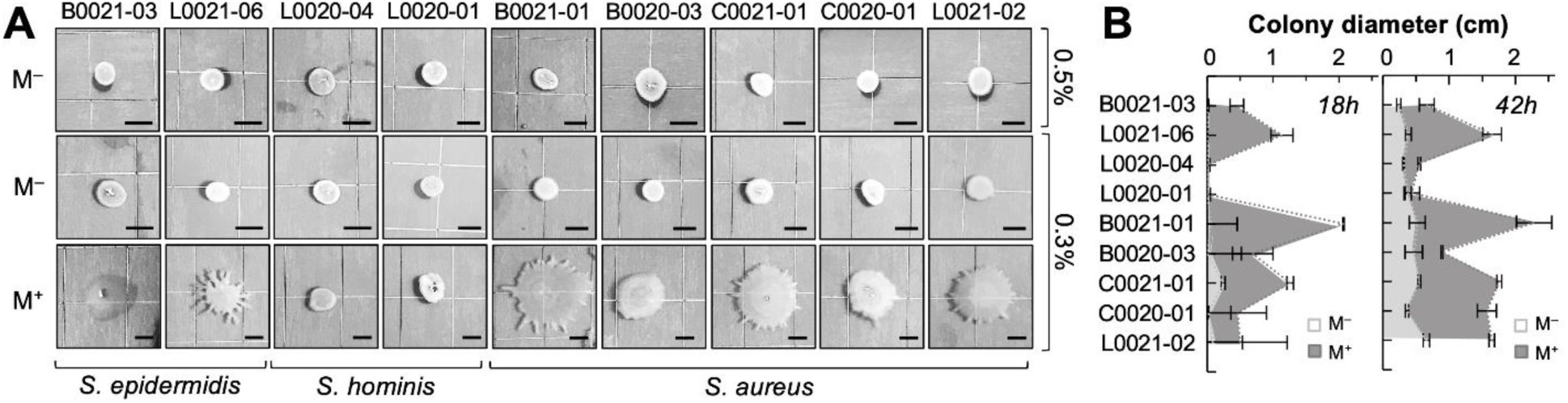
Mucin-induced spreading of (peri)oral *Staphylococcus* colonies. **(A)** Colony morphology of staphylococcal (peri)oral isolates grown on 0.5% (w/v) or 0.3% (w/v) TSA plates with (M^+^) or without (M^−^) 0.4% (w/v) purified porcine gastric mucin. All strains were drop spotted (1 µl TSB, OD_600_ 0.1) on the surface of the plate and allowed to dry for 30 min before incubating at 37°C for 18 h. Scale bar, 0.5 cm. **(B)** Average diameter (in cm) and standard deviation of triplicate colonies for each strain after 18 or 42 h growth on 0.3% (w/v) TSA media with (M^+^) or without (M^−^) purified porcine gastric mucin.

### Relationship between mucin growth and surface dispersal by perioral isolates

Commercial porcine stomach mucin (Type II), a well-characterized gastric and gel-forming mucin, also induced the spreading and dendritic expansion of the *S. aureus* and *S. epidermidis* colonies on motility agar plates (**Fig. 2A**), albeit not as strongly as with the purified mucin (**Fig. 1**). This result is not unexpected given the lower purity of the commercial mucin, which effectively decreases the concentration of glycoproteins in the plates and the viscoelasticity of the medium. The commercial gastric mucin also supported the growth of most strains (**Fig. 2B**). In general, there was a positive correspondence between the ability of the strains to disperse on the mucin plates and to use the glycoproteins as growth substrates. For example, the gastric mucin promoted the rapid expansion of *S. epidermidis* L0021-06 colonies (**Fig. 2A**) while also supporting high yields of planktonic growth (**Fig. 2B**). Furthermore, a mucinase plate assay detected a halo of mucin degradation around most of the spreading colonies, which is also consistent with the use of the glycoproteins and their degradation products as growth substrates (**Fig. 2C**). Conversely, the non-spreading *S. hominis* L0020-04 (**Fig. 2A**) did not grow with mucin (**Fig. 2B**) nor did it secrete mucinases (**Fig. 2C**). Notably, all the *S. aureus* strains were robust spreaders and grew with the mucin, in some cases displaying diauxic growth phases as cells transitioned from growing with the tryptic soy nutrients in the broth to mucin-based growth (**Fig. 2B**, arrows). Additionally, all but one (L0021-02) strain of *S. aureus* had detectable levels of mucinase activity (**Fig. 2C**). The correspondence between surface dispersal and mucin growth by the (peri)oral isolates suggests that the two processes are linked.

**FIG 2:**
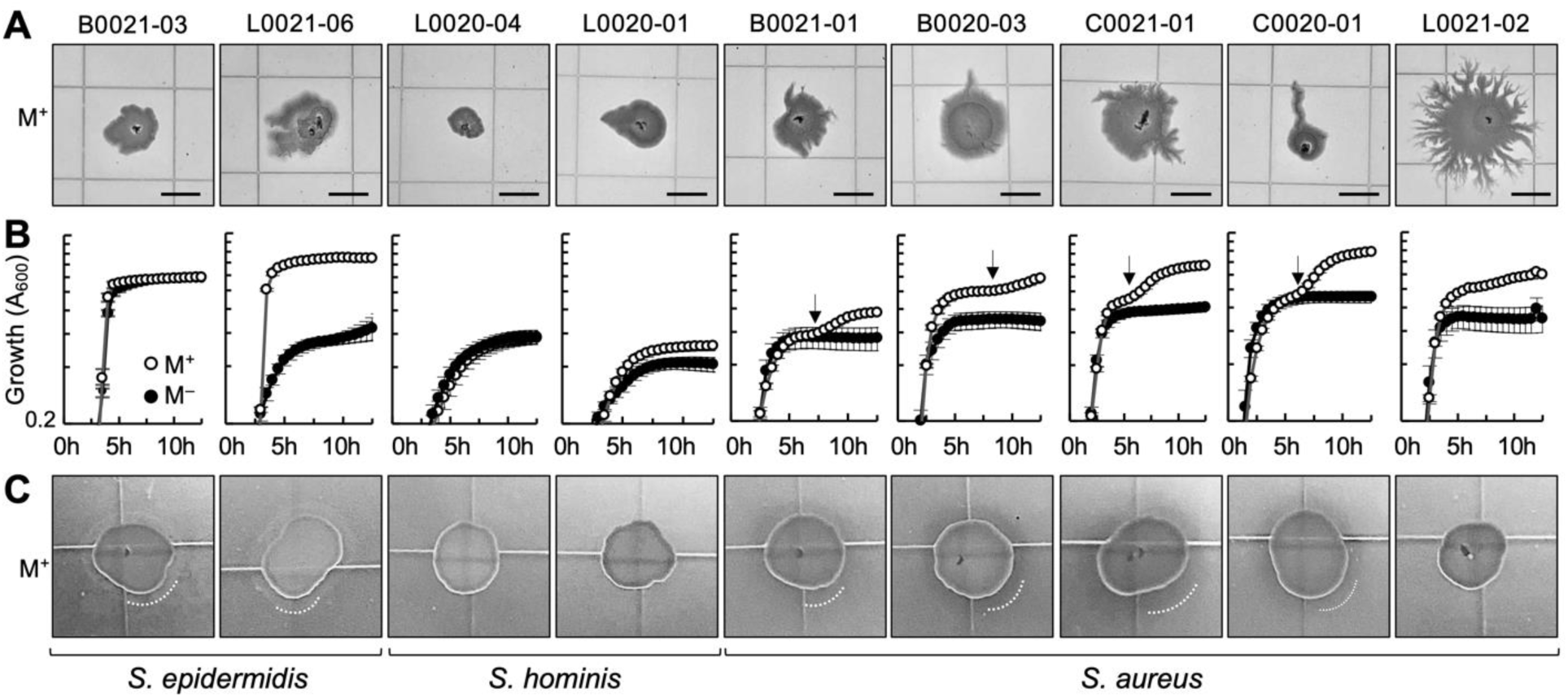
Effect of commercial-grade gastric mucin on dispersal and nutrition of staphylococcal isolates. **(A)** Colony expansion of the (peri)oral staphylococcal isolates on 0.3% (w/v) TSA supplemented with 0.4% (w/v) commercial porcine gastric mucin (M^+^). Incubation was at 37°C for 18 h. Scale bar, 0.5 cm. **(B)** Growth (average optical density, measured as absorbance at 600 nm (A_600_), and standard deviation of 8 replicate cultures) in broth (50% diluted TSB) with (M^+^) or without (M^−^) commercial-grade gastric mucin (0.4%, w/v). Diauxic transitions from nutrients in TSB to mucin growth are marked with an arrow. **(C)** Mucinase activity of colonies grown on 1.5% (w/v) TSA supplemented with 0.5% (w/v) commercial gastric mucin and incubated at 37°C for 24 h. The edges of the zones of mucin degradation, when present, are highlighted in white.

### Conservation of mucin-induced spreading and dendritic expansion phenotypes in laboratory strains closely related to the staphylococcal isolates

We also evaluated whether laboratory derived staphylococcal isolates retained the ability to translocate rapidly in the presence of mucin. As test strains, we selected close relatives of the (peri)oral isolates available in our strain collection: *S. aureus* JE2*, S. epidermidis* RP62a, *Staphylococcus lugdunensis* N920143, and *Staphylococcus haemolyticus* NRS9. The latter two are close relatives of the *S. hominis* isolates used in our studies (**Fig. S1**). We observed in the laboratory strains similar dispersal phenotypes in response to the commercial gastric mucin (**Fig. 3**). For example, *S. lugdunensis* and *S. haemolyticus* shared with the perioral S. *hominis* relatives their inability to spread on the mucin soft-agar (0.3% w/v) surface (**Fig. 3A**), to produce mucinases (**Fig. 3A**), or to grow in broth with mucin (**Fig. 3B**). By contrast, *S. epidermidis* RP62a and *S. aureus* JE2 displayed mucin-induced colony spreading and dendritic expansion (**Fig. 3A**), albeit less robustly than the human, undomesticated isolates (**Fig. 2A**). *S. epidermidis* RP62a, for example, did not spread as robustly as the human isolate L0021-06, although the laboratory strain secreted mucinases (**Fig. 3A**) and grew to high yields with mucin (**Fig. 3B**). Colony spreading and dendritic expansion on mucin plates was also less robust in *S. aureus* JE2 (**Fig. 3A**) than in the closest (peri)oral relatives (**Fig. 2**). Notably, the JE2 strain had mucinase activity (**Fig. 3A**) but did not grow with mucin as a growth substrate in the broth (**Fig. 3B**). Hence, although mucin growth may facilitate rapid surface dispersal, the two processes may not always need to be coupled.

**FIG. 3:**
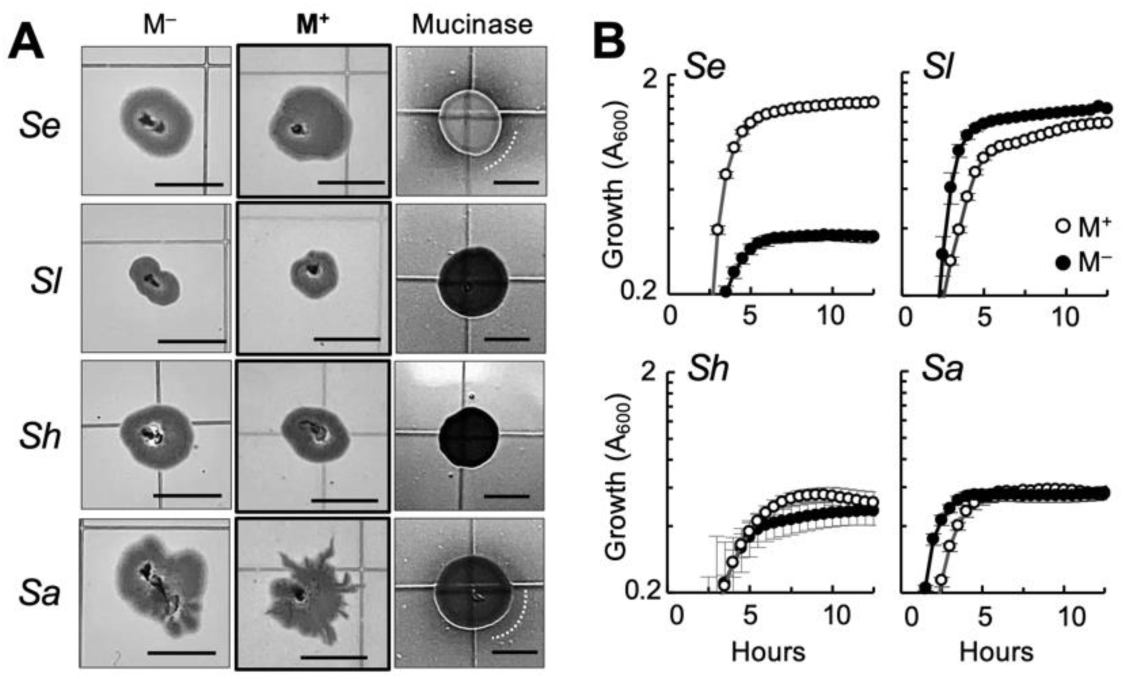
Effect of mucin on dispersal and growth of laboratory relatives of the (peri)oral staphylococcal isolates. **(A)** Morphotype of 18-h colonies of *S. epidermidis* RP62a (*Se*), *S. lugdunensis* N920143 (*Sl*), *S. haemolyticus* NRS9 (*Sh*) and *S. aureus* JE2 (*Sa*) grown at 37°C on 0.3% (w/v) TSA with (M^+^) or without (M^−^) supplementation (0.4% w/v) with commercial gastric mucin or grown with the mucin on 1% (w/v) TSA plates to stain. The panels at right show the mucinase activity of the strains on 1.5% (w/v) TSA supplemented with the mucin (white line shows the edge of the clearing zone, when present). Scale bar, 0.5 cm. **(B)** Average growth (A_600_ of 8 replicates) and standard deviation of the strains in broth (50% TSB) with (M^+^) or without (M^−^) the commercial gastric mucin (0.4%, w/v).

### The surfactant activity and hydrating properties of the mucin modulate staphylococcal colony spreading and dendritic expansion

The lubricating properties of mucins are due, in part, to their surfactant activity (22, 23). To visually confirm this, we airbrushed a fine mist of mineral oil droplets (atomized oil assay) onto the surface of the gastric mucin plates and on control plates without mucin (**Fig. 4A**). The mineral oil droplets remained heterogeneously distorted when applied to the hydrophobic surface of TSA plates without mucin but spontaneously adopted spherical shapes soon after deposition on the mucin plates, an energetically favorable shape associated with surfactant-active surfaces such as after treatment with the anionic surfactant Sodium Dodecyl Sulfate (SDS) detergent (**Fig. 4A**). As the hydrating properties of the mucins also impact lubrication (23, 24), we additionally compared the dispersal of the laboratory staphylococcal strains on the gastric mucin plates to plates containing with the less hydrating submaxillary mucin type (**Fig. 4B**). The commercial porcine stomach (gastric) mucin used in the plate assays has a similar size, net charge, and structural conformation (“dumbbell”-like shape) in aqueous solutions as the bovine submaxillary mucin type (25). However, the uniform distribution of the glycosylated moieties along the gastric glycoprotein increases its hydration compared to the submaxillary type, which concentrates the glycosylation in the center (16, 25). As predicted from their hydrating properties, only the gastric mucin stimulated the growth and rapid expansion of the staphylococcal spreading strains (*S. aureus* JE2 and *S. epidermidis* RP62a) (**Fig. 4**). The stimulatory effect was most significant on the *S. aureus* JE2 colonies, whose diameter at 18 h was already much larger than colonies grown on control plates without mucin (**Fig. 4**). Furthermore, mucin-induced spreading of the JE2 colonies continued for up to 62 h (**Fig. 4**). By contrast, the submaxillary mucin inhibited the growth of most strains on the plates (**Fig. 4**). This is consistent with the reduced hydration of submaxillary mucin hydrogels (25), which reduces water carriage and nutrient mobilization through the semisolid medium.

**Fig. 4:**
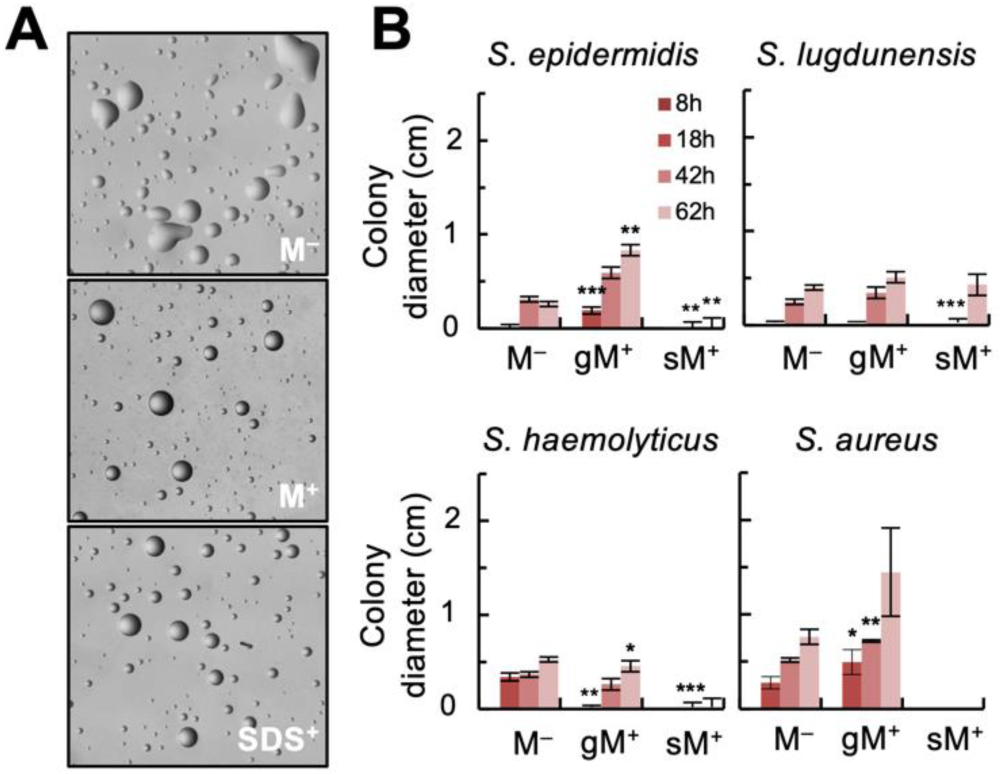
Effect of mucin lubrication (surfactant activity and hydration) on staphylococcal colony growth and expansion. **(A)** Atomized oil assay showing the formation of spherical oil droplets on surfactant-active 0.3% TSA plates containing 0.4% (w/v) of the commercial porcine stomach mucin (M^+^) in reference to plates without mucin (M^−^) or treated with the anionic surfactant SDS (SDS^+^). **(B)** Time-course expansion of colonies (maximum colony diameter, in cm) formed by the laboratory strains of *S. epidermidis* RP62a*, S. lugdunensis* N920143*, S. haemolyticus* NRS9 and *S. aureus* JE2 on 0.3% (w/v) TSA with (M^+^) or without (M^−^) 0.4% (w/v) commercial porcine stomach (gastric, gM) or the less hydrating bovine submaxillary (sM) mucin. The asterisks indicate statistically significant differences (two-tailed, two-sample unequal variance t-test) between the mucin-induced expansion values compared to the M^−^ controls at any given time: ≤0.05 (*), ≤0.005 (**), ≤0.0005 (***).

### Mucin-induced dendritic formation but not colony spreading in *S. aureus* JE2 is regulated by quorum sensing

Leveraging the genetic tractability of *S. aureus* JE2, we used this strain to investigate the molecular mechanisms controlling mucin-induced surface dispersal. Our previous time-course experiment revealed significant colony spreading and dendrite formation of JE2 colonies on gastric mucin plates already at 18 h (**Fig. 3A**). Prolonged incubation (42 h) led to further spreading and consolidation of the dendrites, forming colonies with undulated edges (**Fig. 5A**) and twice the perimeter of colonies grown on the non-mucin controls (**Fig. 5B**). The perimeter of the 42-h colonies thus provided a quantitative method to compare the mucin-induced expansion of the wild-type (WT) colonies to those form by mutants carrying inactivating transposon (Tn) insertions in key components of the *agr* quorum sensing system (*agrA::*Tn*, agrB::*Tn*, agrC::*Tn*, sarA::*Tn) and the *agr*-*sarA* regulated transcriptional regulator *rot* (**Fig. S2**). Inactivation of the *agr* quorum sensing genes (*agrA, agrB* and *agrC*) did not impact the passive spreading of the colonies on mucin plates but prevented their dendritic expansion (colonies had smooth rather than undulated edges) (**Fig. 5A**). As a result, we measured smaller colony perimeters in these mutants compared to the WT (**Fig. 5B**). For example, inactivation of the AgrB autoinducer endopeptidase and export chaperone (26–28) prevented dendritic expansion and reduced the perimeter of the mucin-grown colonies 1.4-fold (p≤0.005). Inactivation of the quorum sensing histidine kinase (AgrC) or the response regulator (AgrA) had a similar effect. Insertional inactivation of the transcriptional enhancer of the *agrACDB* and RNAIII/*hld* operons (*sarA::*Tn mutant) (**Fig. S2**) produced slightly undulated colony morphologies, suggesting a partial defect in dendritic expansion compared to the WT (**Fig. 5A**). Conversely, *sarA::*Tn colonies had larger perimeters than the *agr* mutants but smaller than the WT (p≤0.05) (**Fig. 5B**). This is not unexpected because the inactivation of SarA lowers the levels of secreted PSMs in an *agr*-dependent manner, thereby reducing lubrication (29). It is unlikely that protease hyperproduction in this mutant (29) influenced dispersal significantly, because another mutant with exacerbated protein secretion (*rot::*Tn) grew into colonies that were indistinguishable from WT (**Fig. 5A**). Furthermore, the inactivation of secreted proteases (including aureolysin) in *S. aureus* LAC (ΔESPN) did not significantly affect mucin-induced dispersal, although colony morphotypes and perimeters after 18 h were more variable in the mutant than in the parental strain (**Fig. S3**). Phenotypic variability during dispersal may have resulted from the pleiotropic effects of the ΔESPN mutation on PSM processing and production (29–31), which can reportedly increase the levels of the surfactant active PSMα3 (32). However, we did not observe differences in surfactant activity between the WT and mutant colonies with the atomized oil assay (**Fig. S3**). Moreover, mucinase activity was also similar in the two strains and both were unable to grow with mucin in planktonic cultures (**Fig. S3**). Taken together, these results point to *agr*-regulation of PSM secretion rather than proteases as the primary mechanism for control of mucin-induced dendritic dispersal in *S. aureus*.

**Fig. 5:**
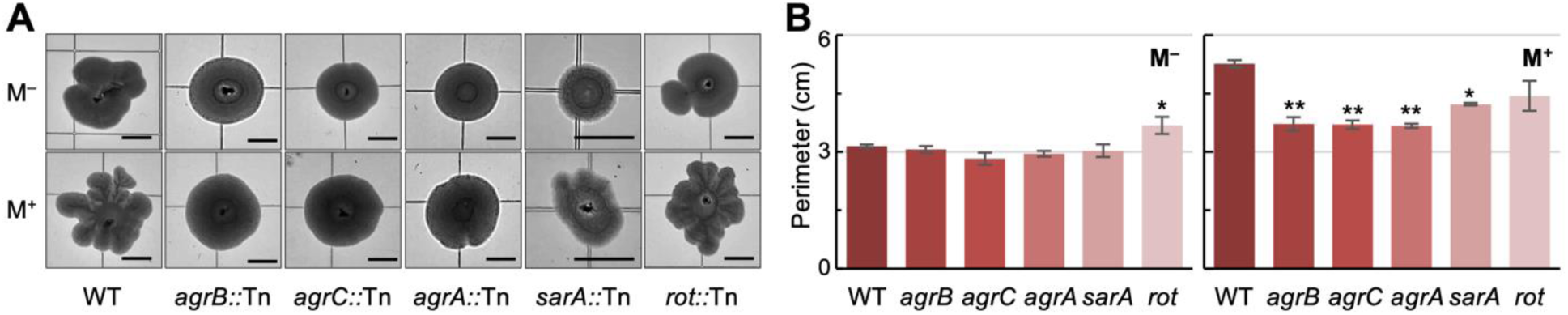
Effect of mucin on the expansion of 42-h colonies of *S. aureus* JE2 and quorum sensing mutants. Morphology (**A**) and perimeter (average and standard deviation of 5-6 replicates from two independent experiments) of 42-h old colonies of *S. aureus* JE2 wild type (WT) and isogenic mutants grown on 0.3% (w/v) TSA plates with (M^+^) or without (M^−^) commercial gastric mucin. The mutant strains carried an inactivating transposon insertion in genes encoding proteins of the quorum sensing regulatory network: AgrB transporter, AgrC histidine kinase for autoinducer sensing, AgrA response regulator, SarA enhancer of AgrA binding, and Rot transcriptional regulator (**Fig. S3**). Statistically significant differences between the mutant and WT colony perimeters (two-tailed, two-sample unequal variance t-test) are marked with asterisks: ≤0.05 (*), ≤0.005 (**).

Further confirming the role of PSMs as facilitators of staphylococcal dendritic expansion, the aerosolized mineral oil assay (33) detected surfactant activity around strains (WT, *sarA::*Tn and *rot::*Tn) with active quorum sensing signaling networks for the endogenous production of PSMs (**Fig. S4**). Moreover, the surfactant-active halo was smaller in the *sarA::*Tn than in the WT (**Fig. S4**), consistent with the lower levels of PSMs secreted by the mutant strain (34). By contrast, we did not detect surfactant-active haloes around PSM-deficient colonies (**Fig. S4**), which included those formed by mutants unable to produce, sense or respond to the autoinducer peptide (*agrA::Tn, agrB::Tn, agrC::Tn*) (35). We also observed a positive correspondence between PSM secretion and mucinase activity in these strains (**Fig. S4**) despite their inability to grow with mucin (**Fig. S5**). As mucin is degraded around the colonies, the viscoelastic properties of the surrounding medium change and PSM lubrication becomes indispensable for cell displacement. Alternatively, PSM lubrication may have increased the solubility of the mucinase enzymes, promoting their diffusion from the colony edge. Regardless of the mechanism, the results suggest that coordinated control of mucin and PSM lubrication is needed for rapid dendritic expansion of *S. aureus* JE2 at high cell densities.

### The surfactant activity of staphylococcal PSMs facilitates dendritic dispersal

The rapid dendritic expansion of colonies produced by (peri)oral isolates of *S. aureus* and *S. epidermidis* on mucin plates (**Fig. 2**) compared to the laboratory strains (**Fig. 3**) suggested greater surfactant activity by PSMs in the human isolates compared to the laboratory relatives. To test this, we used the atomized oil assay to measure the surfactant-active zone (oil dispersion halo) around colonies of representative perioral isolates (*S. aureus* L0021-02, *S. epidermidis* L0021-06 and *S. hominis* L0020-01) and the laboratory strain *S. aureus* JE2 (**Fig. S6**). All but one strain (*S. hominis* L0020-01) exhibited surfactant-active zones around the colonies. The surfactant defect of the *S. hominis* isolate matched well with its inability to expand dendritically on mucin plates (**Fig. 1**) and the reported inability of other *S. hominis* strains to produce lubricating PSMs (14, 36). By contrast, the perioral isolate *S. epidermidis* L0021-06, which spread dendritically on mucin plates (**Fig. 1**), produced large zones of surfactant activity around the colonies (**Fig. S6**). The two *S. aureus* colonies (L0021-02 and JE2) were also surrounded by a surfactant-active halo, albeit smaller than *S. epidermidis* (**Fig. S6**). The halo size in the atomized assay can be influenced by not only the quantity, but also the solubility of surfactant-active molecules secreted by the colonies (33). Hence, *S. aureus* may produce less surfactant active PSMs than *S. epidermidis* and/or the chemistry of the PSMs may be different in these strains. The latter can impact the solubility of the PSMs and their diffusion in the soft-agar medium. This, in turn, reduces the size of the oil dispersal zone around the colonies (**Fig. S6**).

To further investigate the potential correlation of PSM production and dendritic expansion, we sequenced large scaffolds from the genomes of the three representative human isolates and assemble the Illumina short reads to close the genomes. We identified in the sequenced genomes of the robust spreaders (*S. aureus* L0021-02 and *S. epidermidis* L0021-06) all the *psm* genes reported for the laboratory relatives (**Fig. S7**). By contrast, the non-spreading *S. hominis* L0020-01 isolate only encoded *psm* genes of the beta type (**Fig. S7**). The *S. aureus* and *S. epidermidis* genomes also encoded distinct *agr* systems (**Fig. S7**). The genomic analysis is consistent with the divergent quorum sensing networks and PSMs types reported for *S. aureus* and *S. epidermidis* species (37, 38). Species-specific differences in PSM chemistry (including surfactant activity) are therefore expected for these two isolates and are consistent with reports in the literature (28, 37). This is also supported by the higher surfactant activity detected around colonies of *S. epidermidis* L0021-06 with the atomized oil assay, but not with a drop assay (**Fig. S6**), which measures the expansion of droplets of PSM extracts on the hydrophobic surface of Parafilm M (39). The lower solubility of PSMs produced by *S. aureus* L0021-02 reduce the surfactant active zone around the colony in the atomized oil assay. But, once extracted from the colony, a higher surfactant activity was measured for the *S. aureus* L0021-02 PSMs extracts with the drop assay than in any other strain (**Fig. S6**).

### Surfactant-active PSMs stimulate interspecies mucin-dependent dispersal

Despite differences in their *agr* and *psm* systems (**Fig. S7**), PSM-containing supernatant fluids recovered from the robust spreaders *S. aureus* L0021-02 and *S. epidermidis* L0021-06 greatly stimulated the dendritic expansion of the *S. aureus* JE2 colonies (**Fig. 6**). Indeed, control JE2 colonies grown on the mucin plates without supernatants had average perimeters of 2.48 (±0.85) cm after 42 h of incubation at 37°C but were between 1.6 and 4-fold greater after co-incubation with the PSM-containing supernatants. The stimulatory effect of the *S. aureus* L0021-02 supernatant was more pronounced when recovered from stationary phase cultures compared to exponential phase (**Fig. 6**), consistent with the reported accumulation of secreted PSMs in *S. aureus* cultures (10). By contrast, the *S. epidermidis* L0021-06 supernatant collected from exponential phase cultures was already highly stimulatory of JE2 dendritic expansion (**Fig. 6A**), increasing colony perimeters to similar levels as with the stationary phase supernatant (**Fig. 6B**). These results are suggestive of distinct mechanisms for cell density control of PSM production in the *S. epidermidis* and *S. aureus* isolates. Importantly, they demonstrate intra-species and interspecies PSM complementary between the *S. aureus* and *S. epidermidis* spreading strains, which may facilitate co-dispersal.

**Fig. 6:**
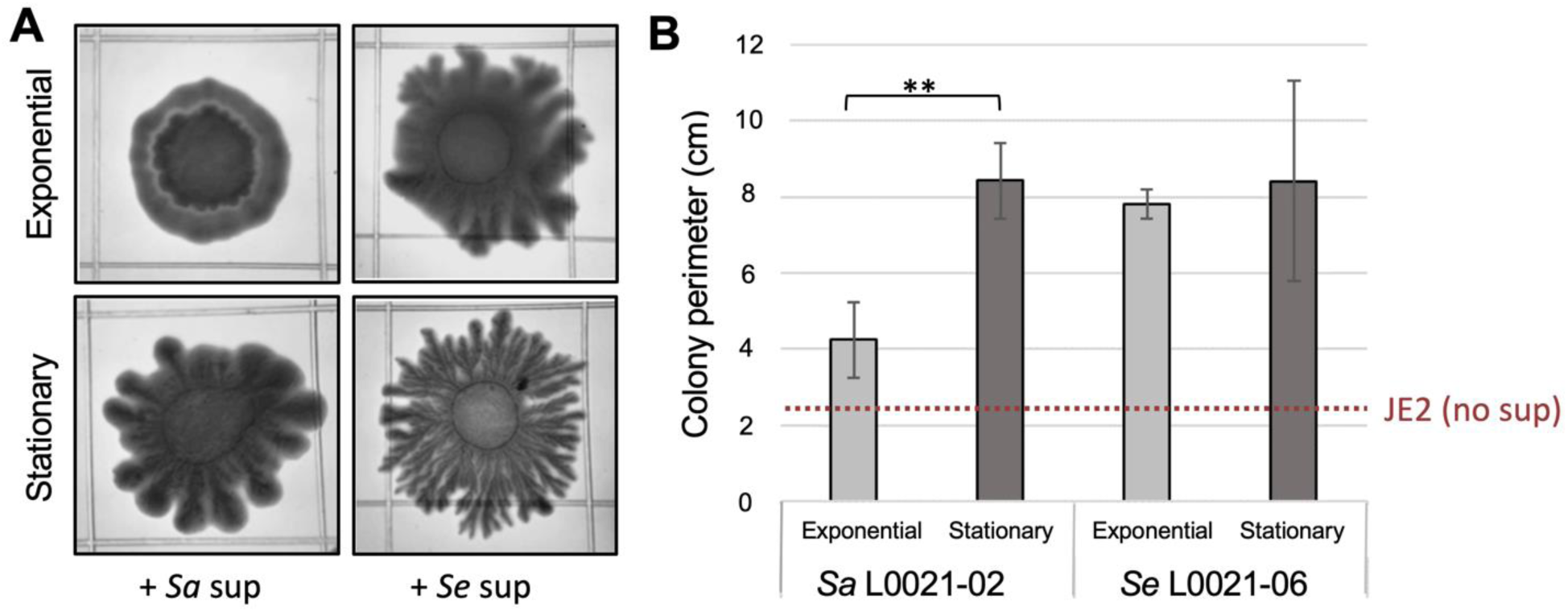
Stimulation of *S. aureus* JE2 dendritic expansion by PSM-containing supernatants from robust perioral spreaders. Supernatant fluids from mid-exponential or stationary phase cultures of *S. aureus* (*Sa*) L0021-02 or *S. epidermidis* (*Sa*) L0021-06 were mixed with *S. aureus* JE2 cells prior to spot plating on TSA 0.3% (w/v agar) plates supplemented with 0.4% (w/v) commercial gastric mucin. The colonies were photographed after 18 h of incubation at 37°C (**A**) and their perimeter measured with the Image J software. The square grid (13 x 13 mm) on the plates is used as a scale. Panel **B** shows the average perimeter of triplicate colonies for each condition in reference to untreated (no supernatant) JE2 colonies (dashed line). Statistically significant differences between the exponential and stationary treatments (two-tailed, two-sample unequal variance t-test) are shown (≤0.05*, ≤0.005**, ≤0.0005***).

In contrast to their stimulatory effect on JE2 colonies, PSM-containing supernatants from robust perioral spreaders (*S. aureus* L0021-02 or *S. epidermidis* L0021-06) did not stimulate dendritic expansion of the perioral *S. hominis* isolate (**Fig. 7A**). The inability of this strain to produce lubricating PSMs (**Fig. S7**) likely prevented interspecies complementarity during colony dispersal. Similarly, the supernatants from the robust spreaders did not stimulate the colony expansion of the isogenic quorum-sensing mutant *agrA::*Tn of *S. aureus* JE2, which is deficient in PSM production, in great contrast to their stimulatory dispersal effect on the parental JE2 strain (**Fig. 7B**). Other combinations, such as JE2 supernatants applied to isogenic mutants with full (*agr* mutants) and/or partial (*sarA::*Tn) defects in PSM production, also failed to chemically rescue the mutant’s dispersal phenotypes (**Fig. S8**). Thus, active quorum sensing networks are likely needed to coordinate PSM production during mucin-induced dendritic dispersal.

**Fig. 7:**
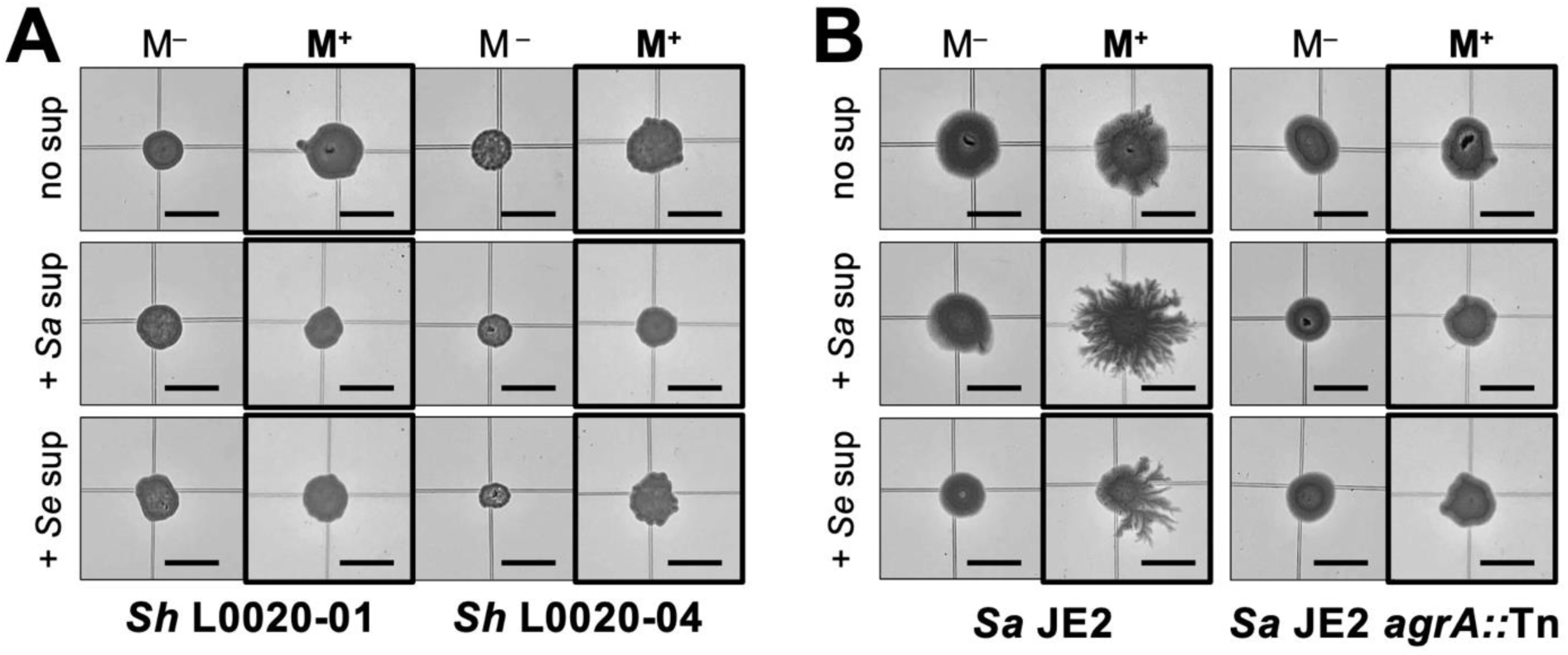
Effect of PSM-containing supernatants from robust perioral spreaders on non-spreading strains. Representative morphotypes of 18-h colonies of non-spreading *S. hominis* perioral isolates L0020-01 and L0020-04 **(A)** and the *S. aureus* JE2 and non-spreading *agrA::*Tn mutant **(B)** on 0.3% TSA without (M^−^) or with (M^+^) porcine stomach mucin treated or not treated with supernatants from the robust spreading perioral isolates *S. aureus* (*Sa*) L0021-02 and *S. epidermidis* (*Se*) L0021-06.

## Discussion

Results from this study demonstrate that the lubricating and hydrating properties of mucins can stimulate passive spreading of staphylococcal colonies and rapid dendritic expansion of cells from the colony periphery in strains (*S. aureus* and *S. epidermidis*) with active quorum sensing (*agr*) systems for controlled secretion of surfactant active PSMs. Surface motile phenotypes have previously been observed in some strains of *S. aureus,* but the reproducibility and biological relevance of these results has been questioned (7). Concerns have been raised, for example, about the highly hydrated agar (0.24-0.34% w/v) conditions that are needed to promote water carriage and passive spreading of *S. aureus* colonies and the effect that drying time (10-30 min) may have on hydration levels and assay reproducibility (8, 9). Dendrites reportedly formed under some of these conditions, but it was unclear whether the phenotype reflected an active mechanism for social dispersal (9). In our study, (peri)oral isolates and close relatives among laboratory strains did not spread or form dendrites on similarly hydrated (0.3% w/v agar), 30 min drying time) surfaces. However, supplementation with lubricating gastric mucins stimulated both the passive spreading and dendritic branching of all the *S. aureus* and *S. epidermidis* colonies (**Fig. 1 and 3**). For these studies, we used undomesticated strains that we previously isolated from the (peri)oral mucosa of healthy young adults (4, 5) and compared their colony phenotypes to those of close relatives among well-characterized laboratory strains such as *S. aureus* JE2 and *S. epidermidis* RP62a. This choice of test strains alleviated concerns previously raised with the use of strains (e.g., *S. aureus* RN4220) that carry genome-wide mutations, including mutations in quorum sensing genes (7). Our plate assays with mucin demonstrated, however, the significant expansion and branching of undomesticated isolates and laboratory strains of *S. aureus* and *S. epidermidis*, unless they carried inactivating mutations in quorum sensing genes (**Fig. 5**). Furthermore, *S. hominis* isolates and closely related strains, which lack quorum sensing networks, did not display the mucin-induced dispersal phenotypes (**Fig. 1 and 3**).

Overnight (18 h) incubation on the mucin plates was sufficient for the motile strains to spread their colonies dendritically, typically increasing the colony diameter 1-2 cm more than in the absence of mucin. Increasing the incubation time (e.g., 42-h assay) enhanced colony expansion and promoted the growth consolidation of the spreading dendrites (**Fig. 1**). We also observed that motile responses of the strains matched well with their ability to grow with the mucin substrate, particularly in the undomesticated strains (**Fig. 2**). This is consistent with the nutritional role that surface dispersal plays in many bacteria (40). In staphylococci, mucin degradation may also help control the rheological properties (e.g., viscosity and elasticity) of the mucus medium to carefully coordinate hydration and lubrications and, thus, coordinate nutritional needs to cell migration. However, nutrition and dispersal on mucin surfaces may be decoupled, as observed in the laboratory strain *S aureus* JE2 (**Fig. 3**).

Using *S. aureus* JE2 as a genetically tractable representative strain, we demonstrated that mucin-induced dendritic expansion was a social behavior regulated by the *agr* quorum sensing system (**Fig. 5**). Surfactants bind mucin glycoproteins and increase the mesh space of the medium, promoting molecular diffusion and water carriage (41). As the rheological properties of the mucin-containing medium change around the edge of PSM-producing colonies, cells can rapidly move away from the colony periphery. This allows the PSM-producing cells to exit areas with high cell density (and potentially, nutrient limitation) and colonize areas where nutrient availability is higher. Consistent with this, the *S. aureus* colonies formed characteristic dendritic projections from the colony after 18 h of incubation, but the dendrites grew and consolidated laterally over time, significantly expanding the diameter and perimeter of the colonies (**Fig. 5**). We observed similar motile behaviors in the *S. epidermidis* isolates (**Fig. 1**) and laboratory relative (**Fig. 3**), consistent with the presence of *agr* quorum systems and *psm* genes in these strains (**Fig. S7**) and their ability to secrete surfactant active PSMs (**Fig. S6**). The rapid dendritic expansion of the *S. epidermidis* strains used in our study contrasted with the “darting” behavior previously reported for this species, which relies on the passive ejection of cell aggregates from the growing colony to slowly spread on the surface (12). Conversely, isolates of *S. hominis* or closely related laboratory strains, which lack most *psm* genes (36), did not spread dendritically. Furthermore, prolonged incubation (42-62 h) was needed to observe significant differences in passive colony spreading between plates supplemented with and without mucin (**Fig. 1 and 4**). These strains were also unable to grow with mucin as a substrate (**Fig. 2 and 3**) or to secrete mucinases for the extracellular degradation of the glycoproteins (**Fig. 3**). These phenotypes correlate well with the ‘dry’ habitats (skin) typically colonized by these staphylococcal species (1) and their lower abundance in the nasal and perioral mucosa compared to robust spreaders such as *S. aureus* and *S. epidermidis* (2–5).

The dispersal potential of *S. aureus* and *S. epidermidis* was influenced by the lubricating and hydrating properties of the mucin glycoproteins. We demonstrated this by comparing the motile phenotype of the laboratory test strains on motility agar supplemented with mucin types (gastric and submaxillary) that share a similar size, net charge, and structural conformation (“dumbbell”-like shape) in aqueous solutions (25), yet influence hydration uniquely due to the different types and distribution of the glycosylated moieties on the protein backbone (16, 25). For example, the gastric mucin has less sialic acid moieties (≤1.2% compared to 9-24% in the submaxillary type) and a more uniform distribution of negative charges along the protein backbone (25). The latter allows the gastric mucin to adopt a conformation that buries the hydrophobic pockets within the glycoprotein and increases its overall hydration (25). This, in turn, facilitates water movement through the gastric mucin hydrogel and reduces frictional forces between the cells and the surface, promoting rapid dispersal. Water movement through the mucin hydrogel also impacts nutrient availability and cell growth. Consistent with this, the more hydrated gastric mucin stimulated the growth and rapid expansion of the staphylococcal colonies while the submaxillary type prevented their growth (**Fig. 4**). The concentration of negative charges in the center of submaxillary mucin glycoproteins (25) may have also increased repulsion forces between the cell and the surface (42), limiting nutrient foraging. Notably, the glycosylation level is lower in the submaxillary (60%) than the gastric (80%) mucin (43), which further reduces the wettability and surface lubrication of the submaxillary mucin hydrogels (16). Given the variable mucin composition of mucosae at each body site (25, 44), mucin-mediated hydration and lubrication may ultimately determine the ability of staphylococci to rapidly disperse and cross-infect specific mucosae. It is perhaps not coincidental that the two staphylococcal species that dispersed rapidly on mucin plates (*S. aureus* and *S. epidermidis*) are also abundant in the mucoid effluents of chronic otitis media patients (45), a condition associated with the hyperproduction of the lubricating and hydrating MUC5B mucin in the middle ear (46). MUC5B and another lubricating mucin, MUC5AC, are also secreted to the respiratory mucosa to modulate the rheology of the mucus layer and promote effective mucociliary clearance in the airways (47). By controlling the hydration and lubrication of the mucus layer (16), mucins can also promote the co-dispersal of staphylococci with other bacteria. For example, mucin lubrication promotes the surface dispersal (“surfing”) of flagellated bacteria such as *P. aeruginosa*, which can carry and co-disperse with *S. aureus* cells (6). Furthermore, through secretion of stimulating factors (including PSMs) at high cell densities (48) and degradation and fermentation of mucin glycoproteins (49), *S. aureus* can potentiate the dispersal and growth of *P. aeruginosa* in the airways, respectively. Unsurprisingly, *S. aureus* and *P. aeruginosa* are often co-isolated from cystic fibrosis airways and other chronic conditions of the respiratory tract (50–53).

Interspecies interactions among staphylococci are also likely to influence their co-dispersal in (peri)oral and respiratory mucosae. In support of this, surfactant active PSM supernatants or colony extracts recovered from the most robust spreading strains of *S. aureus* and *S. epidermidis* stimulated dendritic expansion of the laboratory *S. aureus* JE2 strain greatly (**Fig. 6**). Interspecies complementary thus existed despite notable differences in PSM types and quorum sensing networks in the *S. aureus* and *S. epidermidis* strains (**Fig. S7**). As previously reported for other strains (54), the distinct quorum systems of *S. aureus* (*agr* II) and *S. epidermidis* (*agr* I) influence adaptive responses (biofilm formation and multidrug resistance, respectively) known to impact clinical outcomes (54). Co-dispersal of the two species via quorum sensing complementarity could therefore boost the pathogenic potential of the strains. Given the cross-talk reported for the quorum sensing systems of *S. aureus* and *S. epidermidis* and cross-inhibitory effects of some autoinducer peptides (55, 56), antagonistic effects may also be possible. Antagonistic interactions may also be possible via the secretion of PSM toxins by *S. aureus* (38). Also important is the fact that *S. aureus* secretes PSMs closer to the colony, while *S. epidermidis* extends its surfactant active secretions further away from the cells (**Fig. S6**). *S. aureus* may thus exploit *S. epidermidis* PSMs as “public goods” to promote its own dispersal on mucin surfaces while simultaneously producing antagonistic factors to suppress the growth and co-dispersal of the *S. epidermidis* competitors. Understanding these seemingly contrasting interactions will help predict the dispersal potential of staphylococci from nasal and perioral reservoirs. Importantly, it can inform prophylactic approaches such as bacterial replacement therapies (57) for the treatment of individuals who are most at risk of staphylococcal infections.

## Methods

### Bacterial strains and culture conditions

The bacterial strains used in this study are 9 staphylococcal strains previously isolated by us from buccal, oropharyngeal, and otic secretions collected from 4 healthy young adults (19-32 years old) as a part of a larger study approved by the Michigan State University Biomedical and Physical Health Review Board (IRB # 17-502) (5). The study also included laboratory strains from our culture collection that were most closely related to the (peri)oral isolates (*S. aureus* JE2, *S. epidermidis* RP62a*, S. lugdunensis* N920143, and *S. haemolyticus* NRS9). *S. aureus* JE2 mutants were transposon insertion mutants from the Nebraska TN Library (58), as follows: *agrA::*Tn *erm* NE1532, *agrB::*Tn *erm* NE95, *agrC::*Tn *erm* NE873, *sarA::*Tn *erm* NE1193 and *rot::*Tn *erm* NE386. When indicated, we also included in the study *S. aureus* LAC USA300 strain (59) and its protease-deficient deletion mutant ΔESPN (*S. aureus* LAC strain AH1919) (60). The (peri)oral and laboratory strains were routinely recover from frozen cultures at 37°C in 5 ml of tryptic soy broth (TSB) with gentle agitation (∼200 rpm) overnight, as previously described (4). The overnight culture served as pre-inoculum for growth experiments in TSB or plate assays in tryptic soy medium solidified with various concentrations of agar (TSA).

### DNA extraction, sequencing, genome assembly, and bioinformatic analysis

Strains were grown in 2 ml of TSB at 37°C for 24 h with gentle agitation before harvesting the cells by centrifugation and extracting their genomic DNA, as previously described (4). The DNA was sequenced with the Illumina NextSeq 550 platform at the SeqCenter (formerly Microbial Genome Sequencing Center or MiGS; Pittsburgh, PA) following their standardized protocols. Sequence quality analysis, cleaning/trimming of the Illumina short reads, and de novo assembly into contigs were as described elsewhere (4). We identified the 16S rRNA gene sequences in the contigs with the BAsic Rapid Ribosomal RNA Predictor (Barnap) and deposited the sequences in the GenBank database under individual accession numbers (Supplemental Table 1). To close the genomes of representative isolates (*S. hominis* L0020-01, *S. aureus* L0021-02 and *S. epidermidis* L0021-06) gDNA samples were library prepped using the Rapid PCR barcodking kit (SQK-RPB0004) and sequenced on with Oxford Nanopore Technologies (ONT) Mk1C with R9.4.1 flow cell and ran for 24 h. Sequence quality was assessed with pycoQC v2.5.2 (https://github.com/a-slide/pycoQC), while filtering for read length was performed using filtlong v0.2.1 (https://github.com/rrwick/Filtlong) with its standard command line. Preprocessed short- and long-read sequences were further used in de novo assemblies of staphylococcal isolates with Unicycler v0.4.9 hybrid assembler supported with SPades v3.14.1., following taxonomic ID through MASH2-4. Bioinformatic identification and gene conservation analysis of the *agr* and *psm* genes in the genomes was performed using blastn v2.10.0 (-evalue 1) (61). Visualization of gene conservation among genomes used heatmaps generated with the ggplot package in the R Project software (https://www.r-project.org/).

### Data availability statement

The accession numbers for the 16S rRNA gene sequences of the (peri)oral staphylococci used in this study are listed in Supplemental Table 1. The assembled, closed genomes of *S. hominis* L0020-01, *S. aureus* L0021-02 and *S. epidermidis* L0021-06 were deposited in the GenBank genome database under accession numbers (biosample) JBBBEL000000000 (SAMN40302511), JBBCIO010000000 (SAMN40341640), and JBBCFW000000000 (SAMN40341562) respectively.

### Planktonic growth with mucin

All of the staphylococcal strains used in this study were grown overnight from frozen stocks in 5 ml TSB at 37°C with gentle agitation (∼200 rpm). Overnight cultures (500 µl) were diluted in 5 ml of TSB and grown to mid-exponential phase (OD_600_ ∼0.5-0.6) before a new transfer (500µl inoculum) to 5 ml of TSB. The culture was then grown to stationary phase (OD_600_ ∼0.9-1.0) and diluted in TSB (500 µl in 5 ml). Approximately 18 µl of this cell suspension were added to the wells of a Corning® 96-well clear round bottom TC-treated plate (Corning 3799) and mixed with 162 µl of TSB, 50% TSB (TSB with half the concentration of the tryptic soy) or 50% TSB with 0.4% (w/v) Type II, porcine stomach (gastric) mucin (Sigma Aldrich, M2378). Growth curves for each strain averaged data from 3 biological replicates, each containing data from 8 replicate wells. Plate incubation was at 37°C in a BioTek PowerWave HT plate reader and optical density (OD_630_) was monitored spectrophotometrically every 30 min after gentle agitation for 0.1 s. Each plate included an uninoculated well with the appropriate medium (TSB, 50% TSB or 50% TSB + 0.4% mucin) to use as a blank.

### Motility plate assays

(Peri)oral isolates and laboratory strains were screened for surface dispersal behaviors in motility TSA plates (0.3% w/v agar) with or without 0.4% (w/v) mucin, provided as pure porcine gastric mucin, commercial-grade porcine stomach mucin (Sigma Aldrich, type II, M2378), or commercial-grade bovine submaxillary mucin (Sigma Aldrich, M3895) and prepared for the assays as previously described (79). Pure porcine gastric mucin was kindly donated by Dr. Andrew VanAlst (Dr. Victor DiRita’s laboratory at Michigan State University), who purified the mucin from pig intestines using a previously described protocol (21). When indicated, experiments also included control plates with 0.5% (w/v) TSA (swarming plates). The protocol to culture cells for inoculation on semisolid agar plates is as described elsewhere (4). Briefly, the strains were grown overnight in 5 mL TSB at 37°C with gentle agitation, then back diluted in 1 mL of TSB to a starting OD_600_ of 0.1 to prepare the inoculum for the plate assay. A drop (1 µl) of the cell suspension was spot plated on the agar surface and allowed to absorb at room temperature until dry. The plates were incubated at 37°C for up to 62 h and photographed periodically using an iPhone 11 at 2.4x zoom. We used the measuring tools of ImageJ (62) to calculate the diameter of the central colony (diameter of the original inoculum drop) for each strain and the diameter of all the colonies after incubation. The extent of colony expansion was calculated by subtracting the average diameter of the central colony from the average maximum diameter (in cm) of the expanded colony, including dendrites if present. Dendritic expansion was quantitatively measured as the perimeter of the colony using the ImageJ stock toolkit. For these measurements, we first converted the image to 8-bit type, enhanced the contrast, and converted the image to binary. We then used the “find edges” function in the ImageJ toolbar options to outline the colony edge. If needed, we used the “wand tracing” tool to manually traced any area missed during the automatic edging before measuring the perimeter.

### Mucin Degradation Assay

Selected strains were tested for their ability to secrete mucin degradation (mucinase) enzymes on agar-solidified (1.5% w/v) TSA plates containing 0.5% (w/v) porcine gastric mucin (Type II, Sigma Aldrich). Plate assays with Tn-insertion mutants of *S. aureus* JE2 included erythromycin (10 µg/ml). Overnight cultures (5 µl) were spot plated on the agar surface and allowed to absorb for ∼30 min at room temperature before incubation at 37°C for 24 or 48 h. Mucinase producers produced a halo of mucin degradation around the colonies. The colonies were photographed with an iPhone 11 using backlight and the size of the halo, if present, was measured with the tools in the Image J software.

### Atomized oil plate assay for surfactant detection

We assayed surfactant production from colonies grown in TSA by spraying them with atomized oil, using a modification (4) of a previously described atomized oil assay (33). Briefly, test strains were grown in 5 mL TSB at 37°C with gentle agitation overnight before spot plating 5 µl on agar-solidified (1.5% w/v) TSA plates with and with or without mucin supplementation, as indicated. The drop inoculum was allowed to adsorb at room temperature before incubating the plates at 37°C for 24 h. At the end of the incubation period, we used an airbrush to apply a fine mist of mineral oil droplets on the plate surface. The droplets adopt energetically unfavorable and heterogeneously distorted shapes on the hydrophobic TSA surface but assume more uniform (energetically favorable) and hemispherical shapes around surfactant-producing colonies. This effect raises the droplets, changing the angle of reflection of incident light and producing a visible halo around the colonies.

The atomized oil assay was also used to test the surfactant activity of agar-solidified (1.5% w/v) TSA with or without mucin (0.5% w/v commercial porcine stomach mucin). As positive controls, we surface-treated TSA plates (no mucin) with 5 µl of a 1% (w/v) aqueous solution of sodium dodecyl sulfate (SDS) and incubated at room temperature for ∼30 minutes or until dry before airbrushing the mineral oil. All the plates were photographed immediately after applying the atomized oil using a dissecting scope (Leica MZ6) with a 10X objective and retrofitted with an eyepiece adaptor to an iPhone. The photographs were analyzed with the ImageJ software to calculate the radius of the halo around the colony.

### Drop assay for surfactant activity of PSM colony extracts

We used a previously published protocol (33) to extract PSMs from staphylococcal colonies grown at 37°C on agar-solidified (1.5% w/v) TSA plates for 18 h and rinsed with water prior to extraction. Briefly, we used a sterile wood stick to harvest the colonies from the plates and suspend the biomass in 500 µl of sterile ddH_2_O. The samples were then mixed by vortexing for 10 seconds before assaying the surfactant activity of the PSM extracts with the Parafilm M test (39). This test measures the surfactant activity of a 20 µl drop of the sample as a function of the drop expansion (increased diameter) after deposition for 1 min on the hydrophobic surface of Parafilm M. Distilled water and a 1% (w/v) SDS aqueous solution were used as negative and positive controls, respectively. The drops were photographed with an iPhone and the pictures were analyzed using the ImageJ software to measure the diameter of the drops.

### Isolation and surfactant activity of PSM-containing supernatant fluids

We harvested supernatant fluids from *S. aureus* JE2, *S. aureus* L0021-02, or *S. epidermidis* L0021-06 by centrifugation (Eppendorf 5810R, 3220 g, 10 min) from 10 ml TSB cultures grown overnight (OD_600_ of ∼1; stationary phase). When indicated, supernatants were also harvested from mid-exponential phase cultures. The supernatant samples were then filter sterilized and used to prepare a cell suspension with the test strains. For this, the test strains were grown at 37°C overnight in 5 ml of TSB, diluted in fresh TSB to an OD_600_ of 0.1, and harvested by centrifugation (Eppendorf 5417, 20,817 *g*, 5 min). The cell pellet was then resuspended with equal volumes (1 ml) of filter sterilized supernatant and fresh TSB. The resulting cell suspension was spot plated on 0.3% (w/v) TSA plates with or without 0.4% (w/v) porcine stomach mucin and incubated at 37°C for 8, 18, 42 and 62h before photographing the colonies. When indicated, we used the 18-h plate assays to compare PSM production in stationary versus exponential phase cultures using supernantant fluids recovered from overnight TSB cultures (OD_600_ ∼1, stationary phase) or from cultures grown for 6 h at 37°C (OD_600_ ∼0.8, exponential phase).

## Acknowledgments

The authors would like to thank Dr. Andrew VanAlst from the lab of Dr. Victor DiRita at Michigan State University for supplying purified porcine gastric mucin for this study. We are also grateful to Drs. Batsirai Mabvakure and Heather Blankenship at the Michigan Department of Health and Human Services for assistance with bioinformatic analysis of Illumina sequences. We thank Dr. Alex Horswill from the University of Colorado Anschutz School of Medicine for providing the *S. aureus* LAC strains and acknowledge the *S. aureus* JE2 transposon mutants obtained from the Nebraska Transposon Mutant Library (NTML) Screening Array NR-48501, which are provided by the Network on Antimicrobial Resistance in *Staphylococcus aureus* (NARSA) for distribution by BEI Resources, NIAID, and NIH.

## Authors contributions

All the authors contributed to the study design. KJ and SHV performed the experimental work and, with GR, interpreted the data for this article. GR wrote the paper with partial drafts from KJ and SHV. All authors read and approved the manuscript.

